# Tumor Necrosis Factor Receptor Signaling Modulates Carcinogenesis in a Mouse Model of Breast Cancer

**DOI:** 10.1101/2020.03.12.989335

**Authors:** Ling He, Kruttika Bhat, Sara Duhacheck-Muggy, Angeliki Ioannidis, Le Zhang, Nhan T. Nguyen, Neda A. Moatamed, Frank Pajonk

## Abstract

Proinflammatory conditions have long been associated with mammary carcinogenesis and breast cancer progression. The underlying mechanisms are incompletely understood but signaling of TNFα through its receptors TNFR1 and TNFR2 is a major mediator of inflammation in both, obesity and in the response of tissues to radiation, two known risk factors for the development of breast cancer. Using the MMTV-Wnt1 mouse model for spontaneous breast cancer and knockout mice for TNFR1 and TNFR2 we report that loss of a TNFR2 allele leads to ductal hyperplasia in the mammary gland with increased numbers of mammary epithelial stem cell and terminal endbuds. Furthermore, that loss of one TNFR2 allele increases the incidence of breast cancers in MMTV-Wnt1 mice and results in tumors with a more aggressive phenotype and metastatic potential. The underlying mechanisms include a preferential activation of canonical NF-κB signaling pathways and autocrine production of TNFα. Analysis of the TCGA dataset indicated inferior overall survival for patients with down-regulated TNFR2 expression.

We conclude, that imbalances in TNFR signaling promote the development and progression of breast cancer, indicating that selective agonists of TNFR2 could potentially modulate the risk for breast cancer in high-risk populations.

**Significance Statement:** Over the past four decades the treatment results for patients suffering from breast cancer have constantly improved, leaving breast cancer prevention as an important frontier against the second leading cause of cancer death in the United States. Obesity has become a national health crisis and is a known proinflammatory risk factor for breast cancer. Our study describes a previously unknown aspect of proinflammatory signaling on breast cancer development and progression, thus suggesting novel potential targets to modulate the incidence of the disease.

## Introduction

Since 1975 the incidence of breast cancer in the United States has been steadily rising. For 2019, 268,600 new cases were estimated, making up 15.2 % of all new cancer cases in the U.S. At the same time, 5-year survival rates increased from 75.3 % to 89.9% in 2015 (https://seer.cancer.gov/statfacts/html/breast.html). 41,760 women in the U.S. are predicted to die from breast cancer in 2019 accounting for 6.9% of all cancer deaths in the country.

Major risk factors for the development of breast cancer aside from gender and age are obesity (1) and thoracic radiotherapy during puberty (2). Both obesity and radiotherapy cause systemic or local proinflammatory conditions with elevated TNFα levels in adipose tissue (3) or within normal tissues exposed to radiation (4). TNFα signals through binding to its receptors, TNFR1 (also called p55 or TNFRSF1A) and TNFR2 (p75 or TNFRSF1B), with pathways downstream of TNFR1 and TNFR2 affecting cell death and survival (5).

In this study we tested the hypothesis that TNFα signaling contributes to the development of breast cancer. Using a transgenic mouse model for spontaneous breast cancer development crossed with TNFR1 or TNFR2 knockout animals we demonstrated that loss of one TNFR2 allele not only affects mammary gland development, but also significantly increases the incidence of breast cancer and leads to a more aggressive tumor phenotype.

## Materials and Methods

### Animals

MMTV-Wnt1, p55 and p75 knockout (KO) mice were originally obtained from the Jackson Laboratories (Bar Harbor, ME). All mice were re-derived, bred and maintained in a pathogen-free environment in the American Association of Laboratory Animal Care-accredited Animal Facilities of the Department of Radiation Oncology, University of California (Los Angeles, CA) in accordance to all local and national guidelines for the care of animals. Due to extensive ductal hyperplasia, the female MMTV-Wnt1 transgenic mice cannot lactate, so we crossed the male MMTV-Wnt1 (+) mice with female p55 KO or p75 KO mice to generate heterozygous MMTV-Wnt1-p55^+/−^ and MMTV-Wnt1-p75^+/−^ offspring. Genotyping was performed for the MMTV-Wnt1 transgene. Mice were monitored regularly and tumors were harvested before exceeding humane endpoints.

### Whole mammary gland mounting

The mammary glands from 6-week-old female mice were carefully excised and spread directly onto a glass slide without changing their original *in situ* shape. The tissue was fixed by immersing in Carnoy’s fixative solution (100%EtOH, chloroform, glacial acetic acid; 6:3:1) at 4° C overnight. Glands were hydrated and stained with carmine alum overnight at room temperature. The stained tissues were then dehydrated and cleared in xylene. Images were captured and merged at 4X using a digital microscope (BZ-9000, Keyence, Itasca, IL). Ductal outgrowth was quantified by measurement of the area (mm^2^) covered by the ductal tree in merged images of the mammary gland whole mounts in ImageJ,and the terminal end buds (TEBs) were counted manually.

### Cell lines

The human SUM159PT breast cancer cell line was purchased from Asterand (Detroit, MI). Cells were cultured in log-growth phase in F12 Medium (Invitrogen, Carlsbad, CA) supplemented with 5% fetal bovine serum, penicillin (100 units/ml), streptomycin (100 μg/ml), 5 μg/ml insulin (Eli Lilly, Indianapolis, IN), 0.1% 1M HEPES buffer (4-(2-hydroxyethyl)-1-piperazineethanesulfonic acid, Invitrogen) and 1 μg/ml hydrocortisone (Pfizer, New York, NY). The cells were grown in a humidified incubator at 37°C with 5% CO_2_ and routinely tested for mycoplasma infection (MycoAlert, Lonza). The identity of the cell line was confirmed by DNA fingerprinting (Laragen, Culver City, CA).

ZsGreen-cODC expressing cells were obtained as described in (6). Briefly, cells were infected with a lentiviral vector coding for a fusion protein between the fluorescent protein ZsGreen and the C-terminal degron of murine ornithine decarboxylase. The latter targets ZsGreen to ubiquitin-independent degradation by the 26S proteasome, thus reporting lack of proteasome function through accumulation of ZsGreen-cODC. We previously reported that cancer cell populations lacking proteasome activity are enriched for tumor-initiating cells in glioblastoma, breast cancer and cancer of the head and neck region (6–9) and others have confirmed these findings independently in tumors of the liver, lung, cervix, pancreas, osteosarcoma and colon (10–15). After infection with the lentivirus, cells expressing the ZsGreen-cODC fusion protein were further selected with G418 for 5 days. Successful infection was verified using the proteasome inhibitor MG132 (Sigma, MO).

### Primary breast tumor cell culture

Breast tumor tissues were isolated from MMTV-Wnt1, MMTV-Wnt1 x p55 KO or MMTV-Wnt1 x p75 KO female mice and washed with PBS/1% Penicillin/Streptomycin. For tumor dissociation, Miltenyi C tubes (gentleMACS C tubes, Cat # 130-093-237, Auburn, CA, USA) were preloaded with 100 μl enzyme D, 50 μl enzyme R and 12.5 μl enzyme A provided in the mouse Tumor Dissociation Kit (Cat # 130-096-730, Miltenyi, Auburn, CA, USA) in 2.35 ml of F12 media. The breast tumor tissues were finely chopped with a scalpel and transferred into the C tubes. Tumor tissues were further dissociated by running the m_impTumor_02 program once on the gentleMACS Dissociator (Cat# 130-093-235, Miltenyi). The tubes were next placed in a shaker incubator at 37 °C for 60 minutes. After incubation, the tubes were once again placed in the gentleMACS Dissociator and subjected twice to the same dissociation program as described above. The tubes were briefly centrifuged at 300 x *g* to collect the digested cells at the bottom of the tube. The cells were re-suspended in the enzyme-containing medium and filtered through a 70-μm filter into a 50 ml conical tube. The filter was washed with 10 ml F12 media and centrifuged at 300 x *g* for 7 minutes. The supernatant was aspirated, and the pellet was re-suspended in 2 ml of ACK lysis buffer (Cat # 10-548E, BioWhittaker, Walkersville, MD, USA) to lyse the red blood cells. After 2 minutes, the conical tubes were centrifuged at 300 x *g* for 7 minutes and the supernatant was aspirated. The cell pellet was resuspended in 1-2 ml F12 media and cell numbers were counted using a hemocytometer. Subsequently, the cells were plated onto 100 mm dishes in SUM159PT culture medium supplemented with plasmocin and grown in a humidified incubator at 37°C with 5% CO2.

### Irradiation

Cells were irradiated at room temperature using an experimental X-ray irradiator (Gulmay Medical Inc. Atlanta, GA) at a dose rate of 5.519 Gy/min for the time required to apply a prescribed dose. The X-ray beam was operated at 300 kV and hardened using a 4 mm Be, a 3 mm Al, and a 1.5 mm Cu filter and calibrated using NIST-traceable dosimetry. Corresponding controls were sham irradiated.

### In vitro limiting dilution assays

SUM159PT spheres were dissociated by TrypLE™ and plated in mammosphere media (DMEM-F12, 10 ml/500 ml B27 [Invitrogen], 5 µg/ml bovine insulin [Sigma], 4 µg/ml heparin [Sigma], 20 ng/ml basic fibroblast growth factor [bFGF, Sigma], and 20 ng/ml epidermal growth factor [EGF, Sigma]) into 96-well ultra-low adhesion plates, ranging from 1 to 256 cells per well, along with a single dose of TNFα (1, 10, 50, 100 ng/ml) or vehicle (0.1% BSA). Growth factors (EGF and bFGF) were added every 3 days. The number of spheres formed per well was then counted and expressed as a percentage of the initial number of cells plated.

MMTV-Wnt1, MMTV-Wnt1 x p55 KO and MMTV-Wnt1 x p75 KO mammospheres were cultured from their corresponding monolayer cells. The cells were detached with Trypsin and plated in SUM159PT mammosphere media for sphere formation. The spheres were then dissociated by TrypLE™ and plated into 96-well ultra-low adhesion plates, ranging from 2 to 512 cells per well, along with a single dose of TNFα (1, 10, 100 ng/ml) or vehicle (0.1% BSA) applied. Growth factors (EGF and bFGF) were added every 3 days. The number of spheres formed per well was then counted, expressed as the percentage of the initial number of cells plated, and normalized to the counts in the MMTV-Wnt1 BSA-treated control group.

### Clonogenic survival assay

MMTV-Wnt1, MMTV-Wnt1 x p55 KO and MMTV-Wnt1 x p75 KO monolayer cells were trypsinized and plated in 6-well plates at a density of 2000 cells for MMTV-Wnt1, and 200 cells for both MMTV-Wnt1 x p55 KO and MMTV-Wnt1 x p75 KO per well. The cell culture media was changed every 3 days. After two weeks, the colonies were fixed and stained with 0.1% crystal violet. Colonies consisting of at least 50 cells were counted in each group and presented as percentage of the initial number of cells plated.

### Cell migration assay

Cell migration was quantified using Transwell plates (8 µm pore size; Corning). After 12 hours of serum starvation, 1×10^5^ MMTV-Wnt1, MMTV-Wnt1 x p55 KO or MMTV-Wnt1 x p75 KO primary cells were placed in the insert, with 750 µl cell culture medium containing 5% FBS as the induction medium in the lower chamber. After 16 hours of incubation, unmigrated cells on the upper surface of the insert were removed with a cotton swab. Migrated cells were fixed with formalin and stained with crystal violet. Images were taken randomly on a digital microscope (BZ-9000, Keyence) with 4x magnification and the migrated cell number was counted and quantified by ImageJ.

### Flow Cytometry

Breast cancer-initiating cells (BCICs) were identified based on their low proteasome activity using the ZsGreen-cODC reporter system (8). Five days after irradiation, cells were trypsinized and ZsGreen-cODC expression was assessed by flow cytometry (MACSQuant Analyzer, Miltenyi). Cells were defined as “ZsGreen-cODC positive” if the fluorescence in the FL-1H channel exceeded the level of 99.9% of the parental control cells.

### Immunohistochemistry and immunofluorescence

Formalin-fixed tissue samples were embedded in paraffin and 4 µm sections were stained with hematoxylin and eosin (H&E) using standard protocols. Additional sections were baked for 1 hour in an oven at 65° C, dewaxed in 2 successive Xylene baths for 5 minutes each and then hydrated for 5 minutes each using an alcohol gradient (ethanol 100%, 90%, 70%, 50%, 25%). The slides were incubated in 3% hydrogen peroxide/methanol solution for 10 minutes. Antigen retrieval was performed using Heat Induced Epitope Retrieval in a citrate buffer (10 mM sodium citrate, 0.05% tween20, pH 6) with heating to 95°C in a steamer for 25 minutes. After cooling down, the slides were blocked with 10% goat serum plus 1% BSA at room temperature. for 30 minutes and then incubated with the primary antibody against Ki67 (Abcam, Cat #15580, 1:200), Vimentin (Cell Signaling, Cat #5741S, 1:200), Snail (Abcam, Cat# 85931, 1:400), CD68 (Thermo Fisher Scientific, PA5-32330, 1:100), iNOS (Thermo Fisher Scientific, PA1-036, 1:100) or CD163 (Thermo Fisher Scientific, PA5-78961, 1:500) overnight at 4°C. The next day, the slides were rinsed with PBS and then incubated with ready-to-use IHC detection reagent (Cell signaling, Danvers, MA; 10 µl) at room temperature for 1 h, rinsed, and then incubated with DAB (Cell Signaling) for 3-5 minutes. Tissues were counterstained with Harris modified Hematoxylin (Fisher scientific, Waltham, MA) for 30 seconds, dehydrated via an alcohol gradient (ethanol 25%, 50%, 70%, 90%, 100%) and soaked twice into Xylene. A drop of Premount mounting media (Fisher Scientific) was added on the top of each section before covering up with a coverslip.

For immunofluorescent staining the slides were incubated overnight with a primary antibody against CD31 (Abcam, Cat# 24590, 1:200) at 4°C, rinsed and incubated with secondary antibody Alexa Fluor 594 Goat Anti-mouse immunoglobulin G (IgG) (H/L) antibody (1:1000, Invitrogen) for 60 minutes at room temperature. protected from light, followed with DAPI counter-staining for 5 minutes. The sections were then mounted with Fluoromount^TM^ Aqueous mounting medium and images were taken with a digital microscope (BZ-9000, Keyence).

### Protein extraction and western blotting

The total protein was extracted from FACS sorted ZsGreen-cODC-negative SUM159PT cells treated with TNFα and/or irradiation. Briefly, the cells were lysed in ice-cold RIPA lysis buffer containing proteinase inhibitor (Thermo Fisher Scientific) and phosphatase inhibitor (Thermo Fisher Scientific). The protein concentration in each sample was determined using the BCA protein assay (Thermo Fisher Scientific) and samples were denatured in 4X Laemmli sample buffer containing 10% β-mercaptoethanol for 10 minutes at 95°C. Equal amounts of protein were loaded onto 10% SDS-PAGE gels (1X Stacking buffer - 1.0 M Tris-HCl, 0.1% SDS, pH 6.8, 1X Separating buffer - 1.5 M Tris-HCl, 0.4% SDS, pH 8.8) and were subjected to electrophoresis in 1X Running buffer (12.5 mM Tris-base, 100 mM Glycine, 0.05% SDS), initially at 40 V for 30 minutes followed by 80 V for two hours. Samples were then transferred onto 0.45 μM nitrocellulose membrane (Bio-Rad) for two hours at 80 V. Membranes were blocked in 1X TBST (20 mM Tris-base, 150 mM NaCl, 0.2% Tween-20) containing 5% bovine serum albumin (BSA) for 30 minutes and then washed with 1X TBST. These were then incubated with primary antibodies against Sox2 (Cell Signaling, Cat# 14962S, 1:1000), Oct4 (Cell Signaling, Cat # 2750S, 1:1000), Klf4 (Cell Signaling, Cat # 4038S, 1:1000), cMyc (Cell Signaling, Cat # 5605S, 1:1000), or GAPDH (Abcam, Cat# 9484, 1:1000) in 1X TBST containing 5% BSA overnight at 4°C with gentle rocking. Membranes were then washed three times for 5 minutes each with 1X TBST and incubated with a secondary anti-rabbit horseradish peroxidase-conjugated antibody (Cell Signaling, Cat # 7074S, 1:2000) in 5% BSA for two hours at room temperature with gentle rocking. Membranes were washed again three times for 5 minutes each with 1X TBST. Pierce ECL Plus Western Blotting Substrate (Thermo Fisher Scientific) was added to each membrane and incubated at room temperature for 5 minutes. The blots were then used to expose X-ray films (Agfa X-Ray film, VWR, Cat # 11299-020) in a dark room. GAPDH was used as a loading control.

Nuclear proteins were extracted from MMTV-Wnt1, MMTV-Wnt1 x p55 KO and MMTV-Wnt1 x p75 KO cells. Briefly, the cells were detached and the pellet resuspended in cell lysis buffer with PMSF, DTT and protease inhibitor, incubated on ice for 20 minutes with intermittent tapping and inverting, then vortexed and centrifuged at 12,000 x *g* at 4°C for 10 min. The cytoplasmic supernatant was discarded and the remaining pellet washed and resuspended in nuclear extraction buffer supplemented with PMSF, DTT and protease inhibitor, along with two-time sonication. The samples were incubated on ice for 30 minutes, centrifuged at 12,000 x *g* at 4°C for 15 min and the supernatant containing the nuclear proteins was transferred into fresh tubes and quantified using the BCA protein assay (Thermo Fisher Scientific).

### NF-κB DNA-binding assay

The DNA-binding activity of NF-κB was quantified by enzyme-linked immunosorbent assay using the TransAM*®* NF-κB family activation assay kit (Active Motif North America, Carlsbad, CA) to specifically detect and quantify the DNA-binding capacity of the NF-κB subunits p65, p50, p52 and Rel-B. The assay was performed according to the manufacturer’s protocol and analyzed using a microplate absorbance reader Spectramax M5 (Molecular Devices, San Jose, CA).

### Proteome Profiler^™^ cytokine array

The MMTV-Wnt1, MMTV-Wnt1 x p55 KO and MMTV-Wnt1 x p75 KO primary tumor cells were plated, serum-starved overnight and placed into fresh serum-free medium the next day. Twenty-four hours later, the cell culture medium was collected and centrifuged to remove insoluble materials. Secreted cytokines were measured using the Proteome Profiler Mouse Cytokine Panel A Array Kit (R&D Systems, Minneapolis, MN, USA) according to the manufacturer’s instructions. The kit consists of a nitrocellulose membrane containing 40 different anti-cytokine/chemokine antibodies spotted in duplicate. Briefly, membranes were incubated with blocking buffer at room temperature for 1 hour. Cell supernatants (1 ml) were mixed with a biotinylated detection antibody cocktail at room temperature for 1 hour, and then each was incubated with a membrane overnight at 4°C. The arrays were then washed three times for 10 minutes and subsequently incubated with horseradish peroxidase-conjugated streptavidin for 30 minutes at room temperature. The arrays were exposed to peroxidase substrate (ECL Western blotting detection reagent; Amersham Bioscience). Luminescence was detected using X-ray films, the films were scanned, and signals were quantified using the ImageJ software package. The data were normalized using the internal controls included on each array.

### TCGA data mining and analysis

The Cancer Genome Atlas data set was accessed via the cBioPortal (16, 17) and the TCGA Provisional dataset (captured December 10, 2019) was interrogated. The overall survival data and the expression data (RNA-Seq V2 RSEM Z scores for TCGA Provisional data) for TNFRSF1A and TNFRSF1B, which encode the TNFR1 and TNFR2 receptors respectively, were downloaded. Patients with both gene expression and survival data accessible were used for analysis (N=1078). Kaplan-Meier survival analysis was performed using a Z-score cut-off of 1.0, and the patients were stratified into subgroups with over-expression, under-expression, or normal expression for each receptor. Overall survival times were used to calculate Kaplan-Meier estimates.

### Statistics

All analyses were performed in the GraphPad Prism 8.0 software package. A two-sided Student’s t-tests was used for un-paired comparisons, and a one-way or two-way ANOVA with a post hoc Bonferroni adjustment was used for comparisons between three or more groups. A log-rank test was used to determine the *p*-value for the Kaplan-Meier survival curves. A *p*-value<0.05 was considered as statistically significant. All *in vitro* experiments were performed in at least three independent biological samples.

### Data sharing

All data and methods are included in the manuscript. Tumor-derived cell lines will be made available upon reasonable request.

## Results

### Loss of TNFR2 signaling impacts mammary gland development

In order to explore if TNFα, a major mediator of inflammation, affects the normal mammary gland development via TNFα/TNFR signaling, we utilized knockout mouse strains for the TNF receptors. Using whole mounts of the fourth mammary gland of prepubescent, 6-week old female C57BL/6, TNFR1 (p55) KO or TNFR2 (p75) KO animals (**Figure 1A**), we found a significant increase in the mammary gland area in both, p55 (3.1-fold, *p*-value=0.0001) and p75 (3.6-fold, *p*-value<0.0001) KO animals (**Figure 1B**). Compared to C57BL/6 wild-type animals, p55 KO animals had a significant 1.5-fold (*p*-value=0.0096) increase in the number of terminal end buds (TEBs) (**Figure 1C**). The number of TEBs in p75 KO was even higher and was significantly increased when compared to wild-type animals (1.9-fold, *p*-value<0.0001) or p55 KO animals (1.3-fold, *p*-value=0.0272) (**Figure 1C**). Expression of the MMTV-Wnt1 transgene leads to robust breast cancer formation in C57BL/6 mice (18). A comparison of the mammary gland areas between 6-week old C57BL/6 wild type and MMTV-Wnt1 transgenic animals showed a 1.9-fold (*p*-value=0.0225) increase in MMT-Wnt1 animals, consistent with the published hyperproliferative phenotype of the mammary epithelium in this strain (18). In order to test if p55 or p75 affects tumor formation under a genetic background prone to mammary carcinogenesis we next crossed MMTV-Wnt1 animals with p55 or p75 KO animals leading to animals with a hemizygous loss of the p55 or p75 gene and expression of the MMTV-Wnt1 transgene.

**Figure 1.**
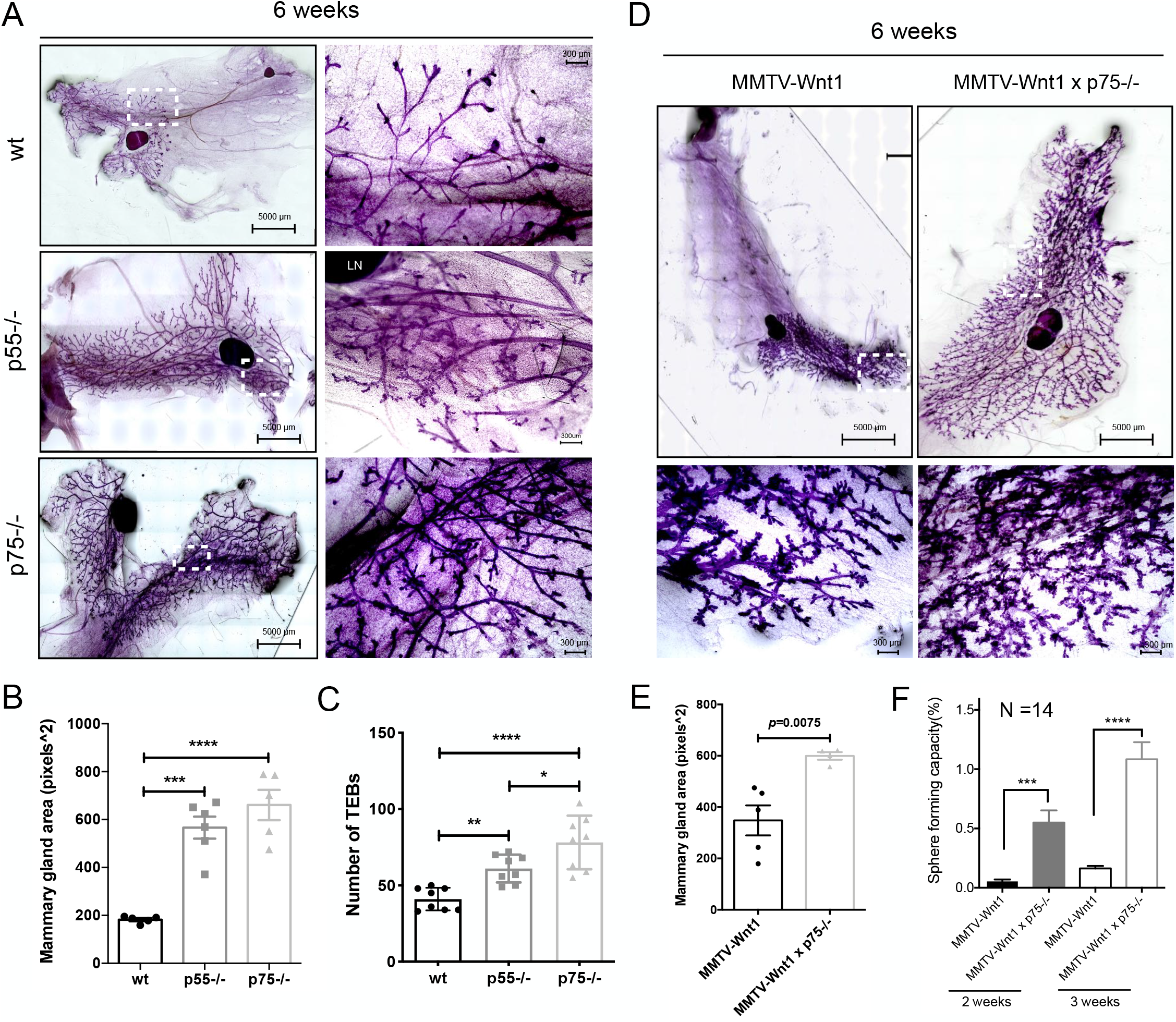
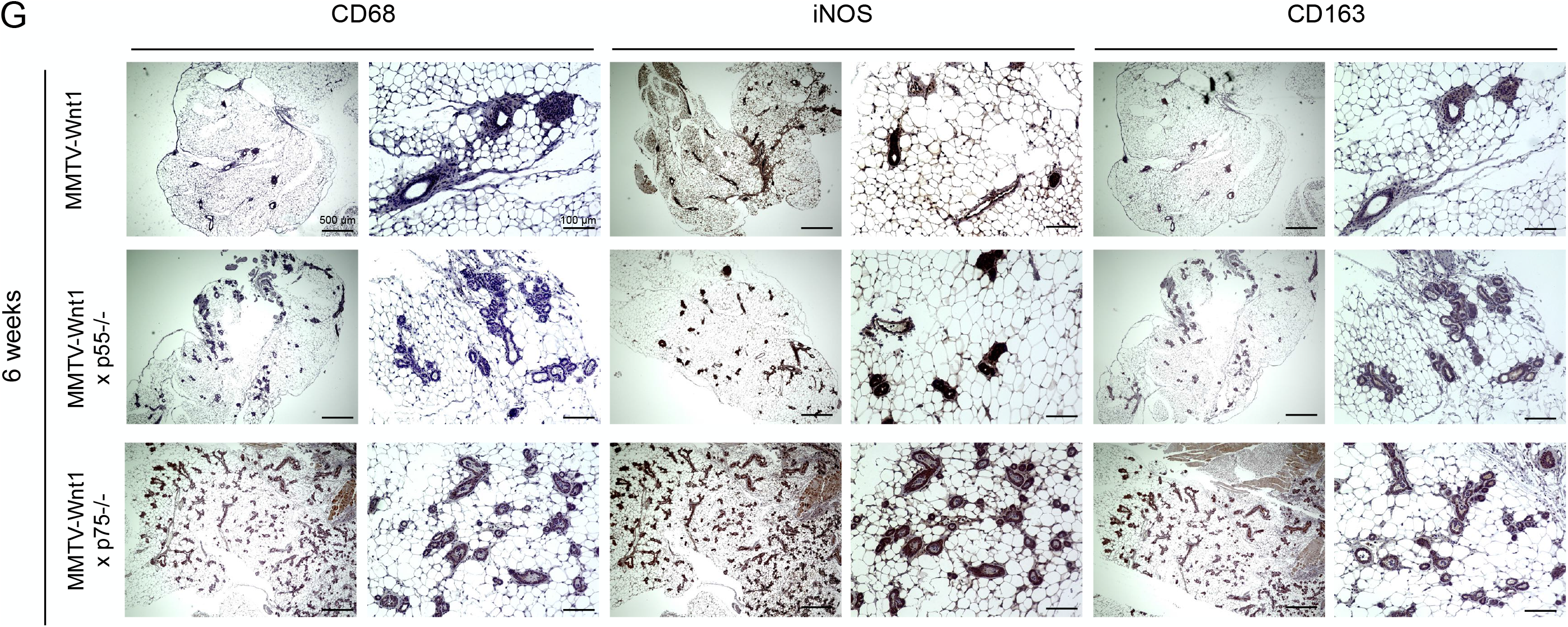
TNFR1/TNFR2 imbalances affect mammary gland development. **(A)** #4 mammary glands were isolated from 6-week-old female C57BL/6 wt, p55 or p75 KO mice, and mounted (LN: lymph node). **(B, C)** Mammary gland ductal outgrowth was quantified by measuring whole mammary gland area using ImageJ and the number of terminal end buds (TEBs) was counted. **(D) Shown here are** whole mount staining of #4 mammary glands dissected from MMTV-Wnt1, MMTV-Wnt1 x p55^−/−^ and MMTV-Wnt1 x p75^−/−^ females. **(E)** Mammary gland ductal outgrowth was quantified by whole mammary gland area using ImageJ. **(F)** Mammary epithelial stem cells were isolated from 6-week-old MMTV-Wnt1 and MMTV-Wnt1 x p75^−/−^ females. Mammosphere forming capacity was evaluated after 2-week and 3-week mammosphere culture. (G) Immunohistochemistry staining of mammary gland sections from 6-week-old MMTV-Wnt1, MMTV-Wnt1 x p55-/- and MMTV-Wnt1 x p75^−/−^ mice using the macrophage markers CD68 (pan-macrophage), iNOS (M1) and CD163 (M2). All experiments have been performed with at least 3 biological independent repeats. *P*-values were calculated using a one-way ANOVA for B and C; un-paired student’s t-test for E and F. * *p*-value<0.05, ** *p*-value<0.01, *** *p*-value<0.001 and **** *p*-value<0.0001.

Mammary gland areas in MMTV-Wnt1 x p75 KO mice were significantly larger than in MMTV-Wnt1 mice (1.7-fold, *p*-value=0.0075) (**Figure 1D/E**) but did not differ from those of p55 KO mice (*p*-value=0.429). The number of TEBs in MMTV1-Wnt1 mice was highly increased over those in wild-type mice and drastically increased in MMTV-Wnt1 x p75 KO mice, which made their quantification impossible (**Figure 1A/D**). In order to test for effects of the TNF receptors on the function of mammary epithelial stem cells we isolated the epithelial cells from the fat pads and subjected them to mammosphere formation assays. Mammary epithelial cells (MECs) from MMTV-Wnt1 x p75 KO mice showed a 10.3-fold increase (*p*-value=0.0003) and a 6.6-fold increase (*p*-value<0.0001) in mammosphere formation at 2 weeks and 3 week in culture respectively, when compared to MECs from MMTV-Wnt1 mice (**Figure 1F**), consistent with increased numbers of TEBs and indicative of increased numbers of mammary epithelial stem cells (19).

Next, we assessed the number of macrophages in mammary glands of 6-week old MMTV-Wnt1, MMTV-Wnt1 x p55 KO, or MMTV-Wnt1 x p75 KO animals. Using the pan-macrophage marker CD68, the M1 marcophage marker iNOS and the M2 marcophage marker CD163 we found elevated number of total, M1 and M2 macrophages associated with the mammary gand ductal system of MMTV-Wnt1 x p75 KO mice when compared to MMTV-Wnt1 or in the MMTV-Wnt1 x p55 KO animals (**Figure 1G**).

### Loss of one TNFR2 allele accelerates breast cancer development in female MMTV-Wnt1 transgenic mice

Next, we assessed the tumor incidences in MMTV-Wnt1 mice and MMTV-Wnt1 mice crossed with p55 and p75 KO animals. Half of the female MMTV-Wnt1 mice developed mammary tumors over the course of two years, which was consistent with tumor incidences reported for this strain in the literature (18). MMTV-Wnt1 x p55 KO mice showed a slightly higher incidence of mammary tumors but this difference was not statistically significant (**Figure 2A**). However, in MMTV-Wnt1 x p75 KO mice mammary tumors developed more rapidly with all animals showing breast tumors within the first year (*p*-value=0.0001) (**Figure 2A**). Histologically, tumors in MMTV-Wnt1 and MMTV-Wnt1 x p55 KO mice were grade 2 breast cancers while tumors in MMTV-Wnt1 x p75 KO mice were less differentiated and predominantly grade 3 carcinomas (**Figure 2B**). Furthermore, tumors from MMTV-Wnt1 x p75 KO mice showed higher degrees of vascularization, with more abundant CD31 positive staining (**Figure 2C**), as well as the epithelial-mesenchymal transition (EMT) markers Vimentin and Snail (**Figure 2D**), indicating the greater metastatic potential in MMTV-Wnt1 x p75 KO tumors. Furthermore, the Ki67 staining showed a trend for increased numbers of proliferating cells in MMTV-Wnt1 x p75 KO tumors (**Figure 2E/F**).

**Figure 2.**
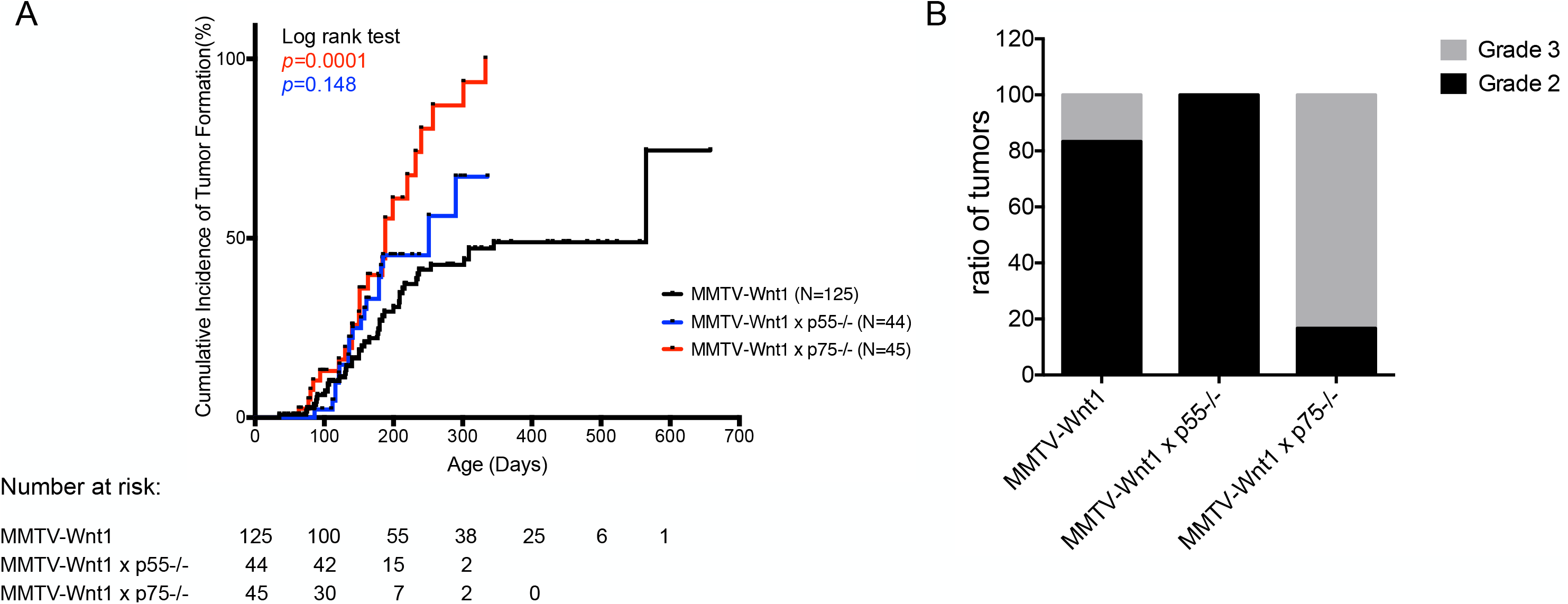
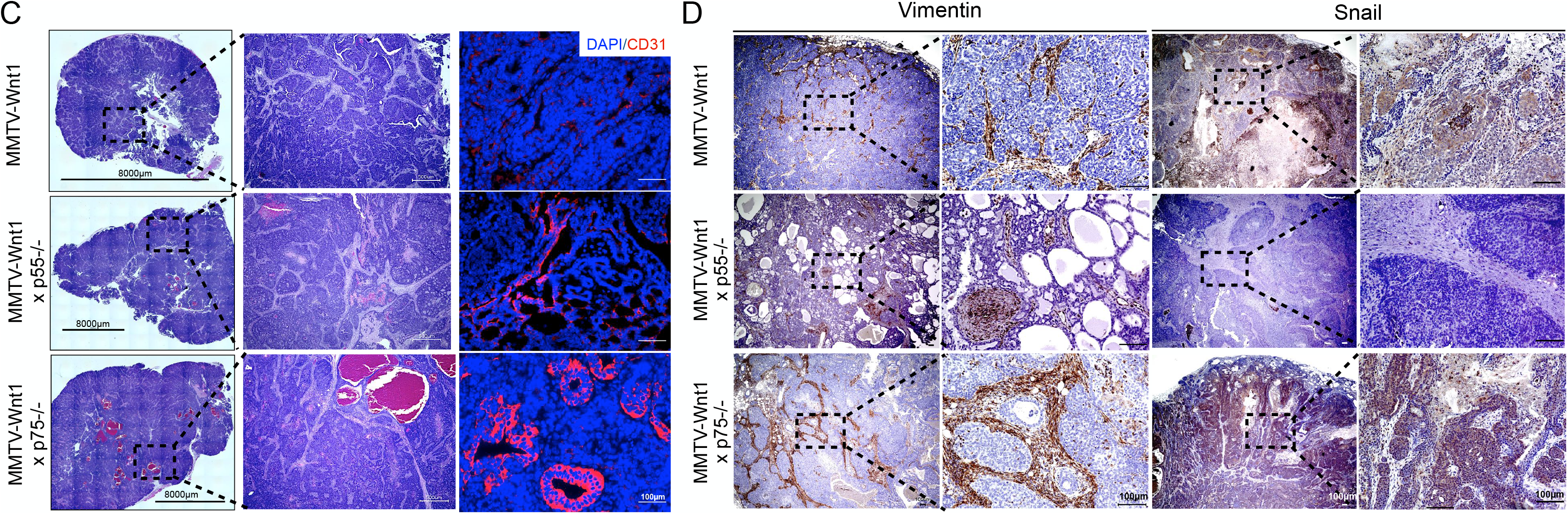
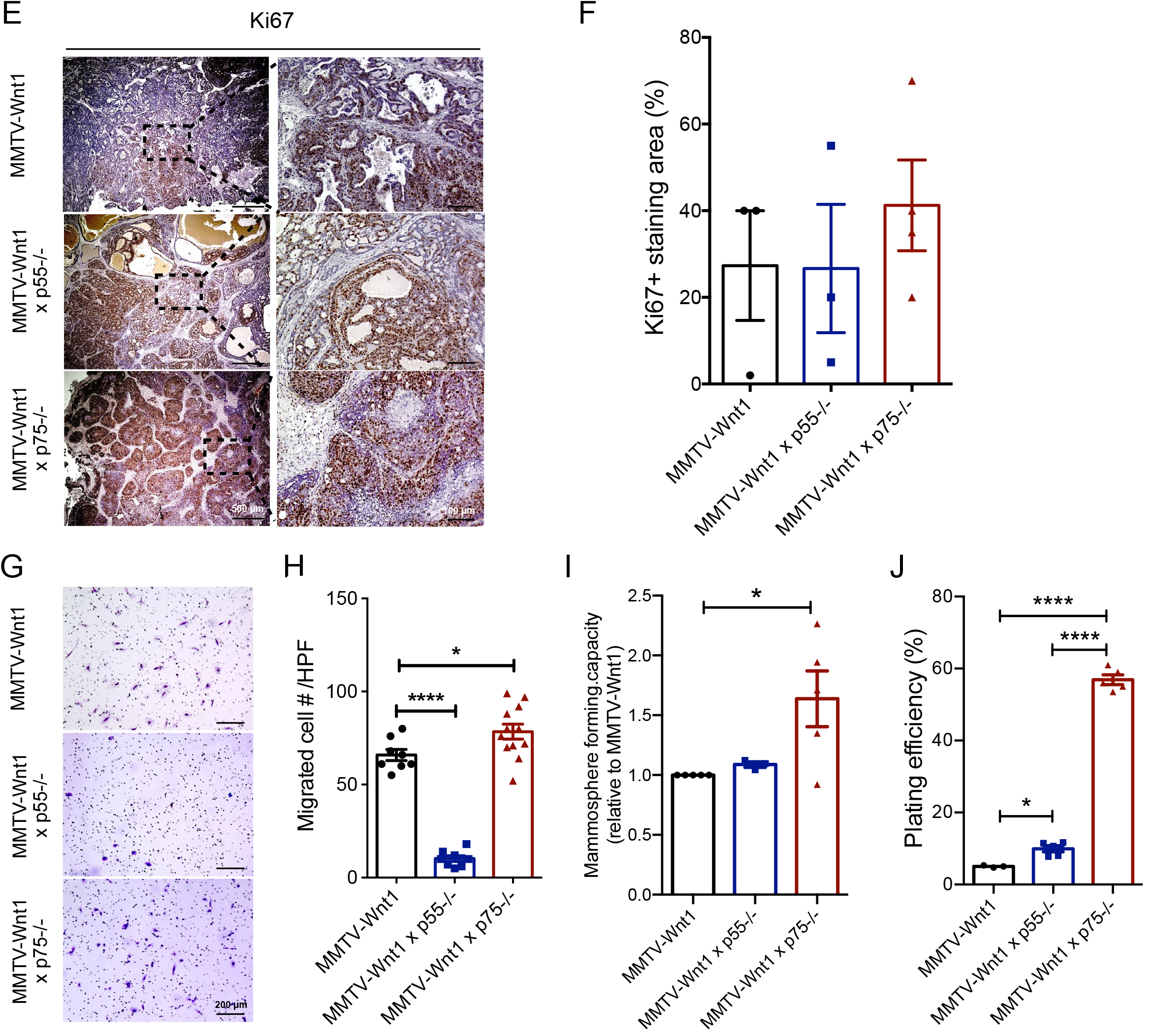
Loss of one allele of TNFR2 increases BCICs and promotes tumor vasculature with greater metastatic potential. **(A)** Cumulative incidence curves for breast cancers in MMTV-Wnt1, MMTV-Wnt1 x p55^−/−^ and MMTV-Wnt1 x p75^−/−^ females. Breast tumors were observed and measured until they reached the study endpoint. Log-rank (Mantel-Cox) test for comparisons of Kaplan-Meier survival curves indicated significant difference between MMTV-Wnt1 x p75^−/−^ and MMTV-Wnt1 mice (*p*-value=0.0001). **(B)** Tumors were harvested, fixed and embedded in paraffin and cut into 4 µm sections. H&E staining was performed and the stained slides were graded blind by a clinical pathologist. **(C)** Representative gross view and the CD31 (PECAM-1) immunofluorescent staining of the tumors from MMTV-Wnt1, MMTV-Wnt1 x p55^−/−^ and MMTV-Wnt1 x p75^−/−^ mice. **(D)** Immunohistochemistry staining of Vimentin and Snail (EMT markers) for the MMTV-Wnt1, MMTV-Wnt1 x p55^−/−^ and MMTV-Wnt1 x p75^−/−^ tumor sections. **(E, F)** Immunohistochemistry staining of Ki67 (proliferating marker) for the MMTV-Wnt1, MMTV-Wnt1 x p55^−/−^ and MMTV-Wnt1 x p75^−/−^ tumor sections, with the slides graded by clinical pathologist for the positive staining area. **(G, H)** Migration assay of primary tumor cells isolated from MMTV-Wnt1, MMTV-Wnt1 x p55^−/−^ and MMTV-Wnt1 x p75^−/−^ mice. The migrated cell numbers were counted and quantified by ImageJ. **(I)** Mammosphere forming capacity of primary tumor spheres isolated from MMTV-Wnt1, MMTV-Wnt1 x p55^−/−^ and MMTV-Wnt1 x p75^−/−^ mice. The number of spheres formed was counted and calculated as a ratio to the initial number of cells plated, and then normalized to the MMTV-Wnt1 value. **(J)** Clonogenic assay of primary tumor cells extracted from MMTV-Wnt1, MMTV-Wnt1 x p55^−/−^ and MMTV-Wnt1 x p75^−/−^ mice. The colony number was counted and presented as the percentage relative to the initial number of cells plated. All experiments have been performed with at least 3 biological independent repeats. *P*-values were calculated using one-way ANOVA for H-J; Log-rank (Mantel-Cox) test for A. * *p*-value<0.05 and **** *p*-value<0.0001.

Next, we validated the more aggressive phenotype of tumors in MMTV-Wnt1 x p75 KO mice using a functional trans-well assay. A comparison of tumor cells from breast cancers in MMTV-Wnt1, MMTV-Wnt1 x p55 KO or MMTV-Wnt1 x p75 KO mice revealed higher migration/invasion capacity in lines established from MMTV-Wnt1 x p75 KO breast cancers (**Figure 2G/H**). Furthermore, cells derived from MMTV-Wnt1 x p75 KO tumors exhibited increased mammosphere formation (**Figure 2I**) and increased clonogenic plating efficiency (**Figure 2J**) when compared to both MMTV-Wnt1 and MMTV-Wnt1 x p55 KO cell lines, thus indicating higher numbers of breast cancer-initiating cells (BCICs) and clonogenic cells, respectively.

### Loss of one TNFR2 allele activates canonical NF-κB signaling

To gain insight into the mechanisms of how knockout of one p75 allele affects mammary gland development and breast carcinogenesis, we next compared the DNA-binding activity of NF-κB family subunits among the primary tumor cell lines using an ELISA-based assay. Nuclear extracts from MMTV-Wnt1 x p75 KO cell lines exhibited significantly higher activation of the p50 NF-κB subunit than MMTV-Wnt1 cell lines, while the MMTV-Wnt1 x p55 KO cell lines had higher activity of the p52 NF-κB subunit and reduced activity of p65 and p50 subunits (**Figure 3A**). Together this suggested preferential signaling through the canonical NF-κB signaling pathway in MMTV-Wnt1 x p75 KO cell lines.

**Figure 3.**
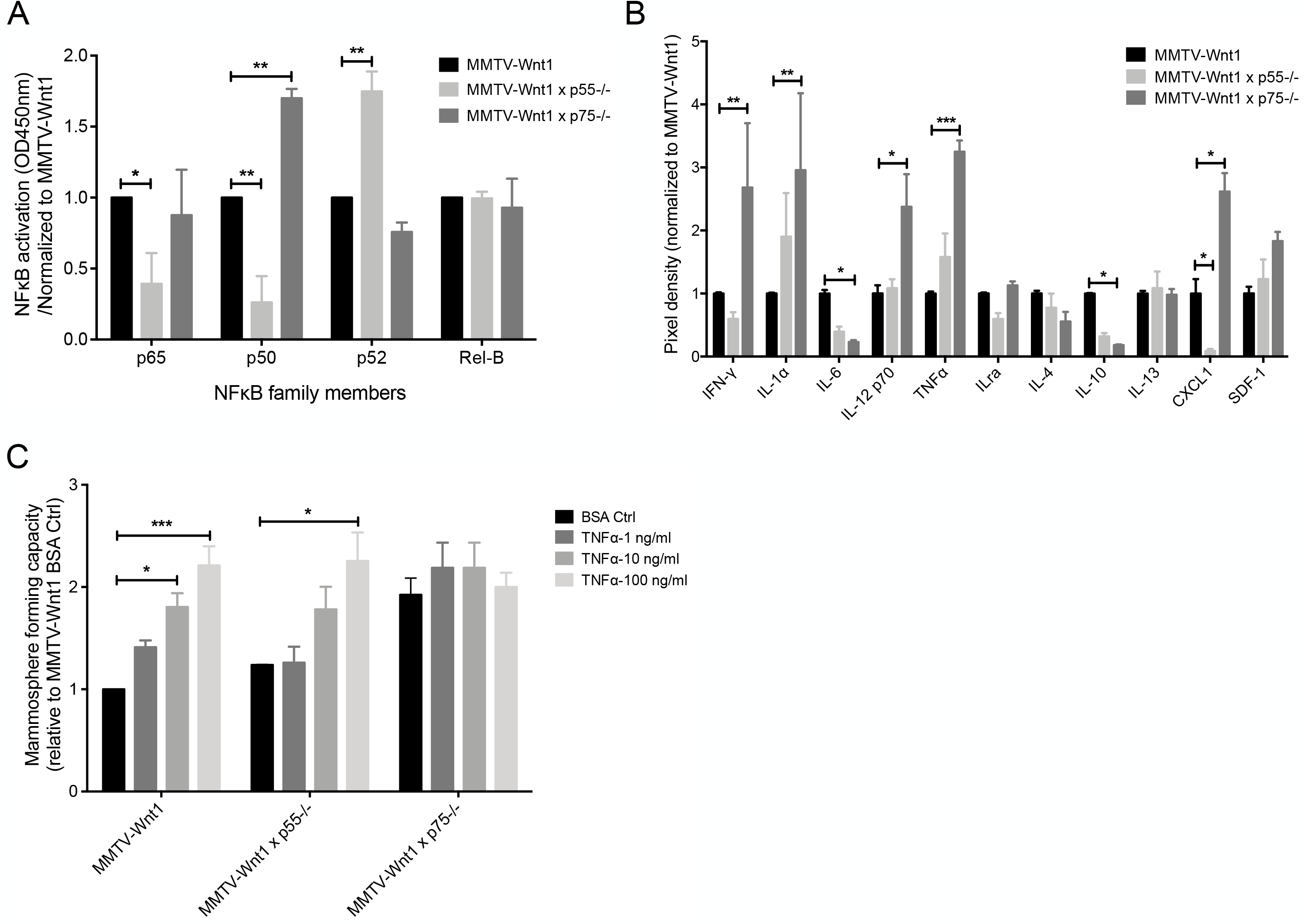
Loss of one TNFR2 allele activates p50/NF-κB transcriptional activity with more abundant secretion of pro-inflammatory cytokines. **(A)** Nuclear extracts prepared from MMTV-Wnt1, MMTV-Wnt1 x p55^−/−^ and MMTV-Wnt1 x p75^−/−^ primary tumor cells were assayed at 5 µg/well for p65, p50, p52 and Rel-B activity using the Trans AM NF-κB Family Kit. The absorbance was normalized to each subunit positive values and subsequently normalization to MMTV-Wnt1 values. **(B)** Proteome profiler cytokine array for nuclear extracts from MMTV-Wnt1, MMTV-Wnt1 x p55^−/−^ and MMTV-Wnt1 x p75^−/−^ primary tumor cell lines. Pixel density was analyzed and normalized to MMTV-Wnt1 by ImageJ. **(C)** Mammosphere forming capacity of primary tumor spheres isolated from MMTV-Wnt1, MMTV-Wnt1 x p55^−/−^ and MMTV-Wnt1 x p75^−/−^ mice upon a single dose of TNFα (1, 10, 100 ng/ml) or vehicle (0.1% BSA) treatment. The number of spheres formed was counted, calculated as a ratio to the initial number of cells plated, and then normalized to the MMTV-Wnt1 BSA control value. All experiments have been performed with at least 3 biological independent repeats. *P* values were calculated using two-way ANOVA. * *p*-value<0.05, ** *p*-value<0.01 and *** *p*-value<0.001.

To further explore the activation of the canonical NF-κB signaling pathway in MMTV-Wnt1 x p75 KO cell lines, we next assessed its pro-inflammatory downstream targets (20, 21) using cytokine arrays. Compared to MMTV-Wnt1-derived lines, MMTV-Wnt1 x p75 KO cell lines showed more abundant secretion of pro-inflammatory cytokines IFN-γ (2.68-fold, *p*-value=0.0097), IL-1α (2.96-fold, *p*-value=0.0023), TNFα (3.25-fold, *p*-value=0.0004) and IL-12 p70 (2.37-fold, *p*-value=0.0393), while the anti-inflammatory cytokine IL-10 (0.19-fold, *p*-value=0.0409) was significantly diminished (**Figure 3B**). Consistent with increased migratory capacity and increased expression of EMT markers, the level of the chemotaxis-related chemokine CXCL1 was increased in MMTV-Wnt1 x p75 KO cell lines (2.62-fold, *p*-value=0.0133, **Figure 3B**).

Finding 3-fold increase of tumor-derived TNFα secretion in MMTV-Wnt1 x p75 KO cell lines, we next explored the effects of exogenous TNFα ligand on self-renewal capacity of MMTV-Wnt1, MMTV-Wnt1 x p55 KO and MMTV-Wnt1 x p75 KO tumor cell lines. Using mammosphere formation assay, we observed a significant increase in mammosphere formation in response to TNFα treatment in MMTV-Wnt1 and MMTV-Wnt1 x p55 KO cell lines, suggesting TNFα could promote self-renewal capacity in breast cancer cells (**Figure 3C**). However, this effect of TNFα was not notable in MMTV-Wnt1 x p75 KO cell lines.

### TNFα-induces phenotype conversion in human triple-negative breast cancer cells

Next, we sought to test the effects of TNFα on SUM159PT human triple-negative breast cancer cells. Treatment with TNFα caused a significant increase in mammosphere formation suggesting that self-renewal in breast cancer cells is regulated by TNFα (**Figure 4A/B**). This effect of TNFα extended to an increased plating efficiency in a classic clonogenic survival assay (**Figure 4C**). We had previously reported that irradiation of non-tumorigenic breast cancer cells led to the induction of a BCIC phenotype (7). Using a reporter system that marks BCICs through accumulation of the fluorescent protein ZsGreen (6, 8, 22) we removed preexisting BCICs by FACS and irradiated the remaining non-BCICs with 0, 4 or Gy in the presence or absence of TNFα. After 5 days in culture, we observed a radiation-induced phenotype conversion of non-BCICs into induced BCICs and an amplification of this effect in the presence of TNFα (**Figure 4D**). This phenotype conversion correlated with the induction of the Yamanaka transcription factors Oct4, Sox2 and Klf4 (**Figure 4E**), which can be used to reprogram somatic cells into induced pluripotent stem cells (iPSCs) (23).

**Figure 4.**
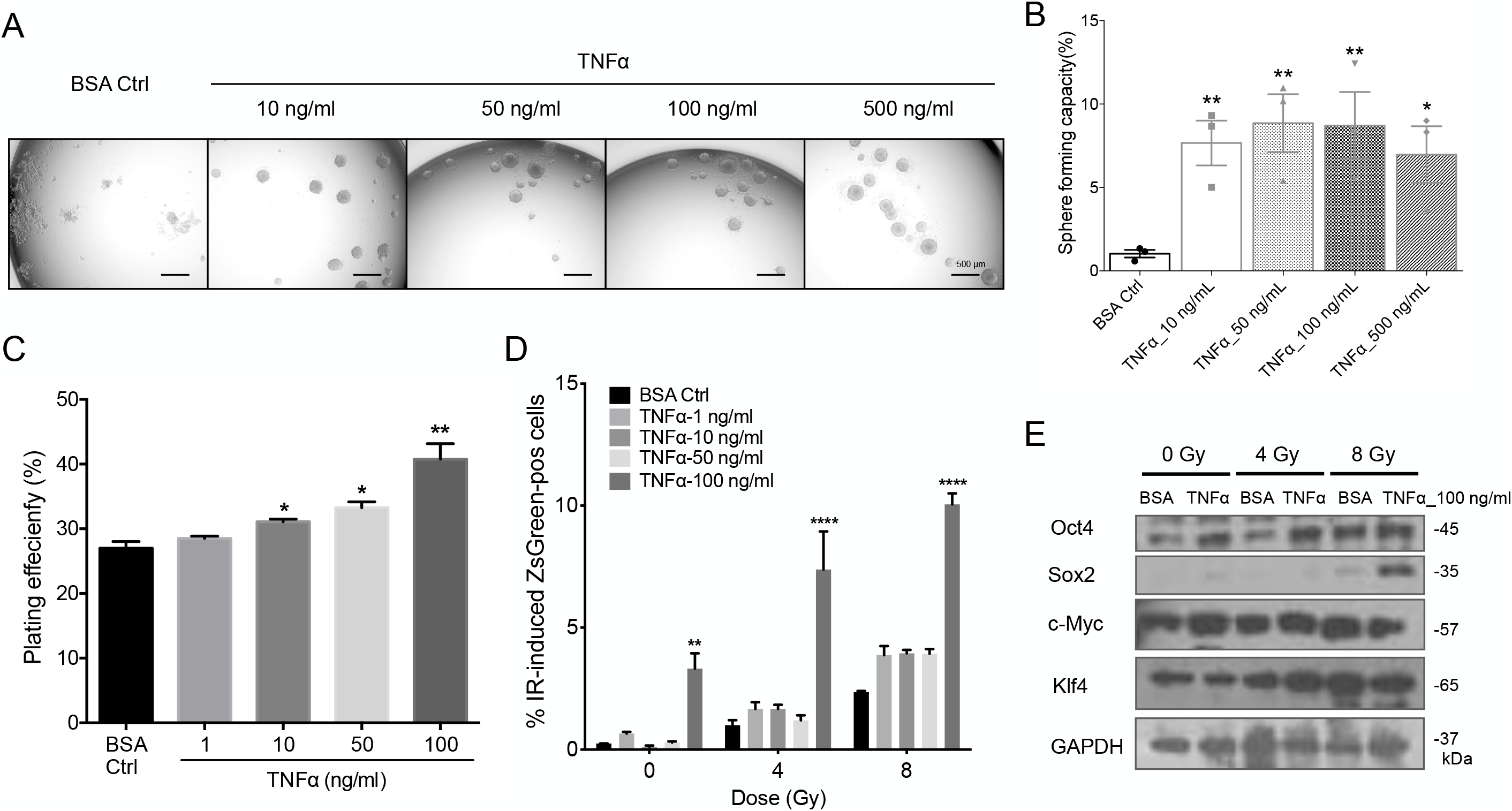
TNFα increases breast cancer initiating cells and induces phenotype conversion in triple negative breast cancer cells. **(A, B)** Mammosphere formation assay was performed using SUM159PT mammospheres plated in a 96-well plate and treated with different concentrations of TNFα (10, 50, 100, 500 ng/ml) or vehicle (0.1% BSA). The number of mammospheres formed in each condition was counted and normalized against the vehicle control. Representative images of mammospheres in each condition (A) with the percentage of mammospheres formed (B) quantified. **(C)** The SUM159PT cells were plated at a density of 200 cells per well in a 6-well plate, then treated with a single dose of TNFα (1, 10, 50, 100 ng/ml) or vehicle (0.1% BSA) and cultured for 14 days. The resulting data was presented as plating efficiency with the percentage of colonies formed. **(D)** Sorted ZsGreen-cODC-negative SUM159PT ZsGreen-cODC vector expressing cells were plated at a density of 50,000 cells per well in a 6-well plate and the following day were pre-treated with either with TNFα (1, 10, 50, 100 ng/ml) or vehicle (0.1% BSA) one hour before irradiation at a single dose of 0, 4 or 8 Gy. Five days later, the cells were trypsinized and analyzed for ZsGreen-cODC-positive population by flow cytometry, with non-infected parental SUM159PT cells used as controls. **(E)** Sorted ZsGreen-cODC-negative SUM159PT cells were plated and pre-treated with 100 ng/ml TNFα or vehicle (0.1% BSA) one hour before irradiation at a single dose of 0 or 4 or 8 Gy. The proteins were extracted five days later and were subjected to western blotting. The blots were analyzed for Oct4, Sox2, c-Myc, Klf4 and GAPDH, with GAPDH as the loading control. All experiments have been performed with at least 3 biological independent repeats. *P*-values were calculated using one-way ANOVA for B and C; two-way ANOVA for D. * *p*-value<0.05, ** *p*-value<0.01 and **** *p*-value<0.0001.

### Down-regulated TNFR2 expression is associated with decreased overall survival in breast cancer patients

To further investigate whether TNFR expression is associated with the clinical outcome of breast cancer patients, we analyzed overall survival (OS) data from 1078 breast cancer patients in the TCGA Provisional dataset, stratified into subgroups with up-, down-regulated or unchanged expression of TNFR1 and 2 using a Z-score cut-off of 1.0. The median survival of patients in the subgroup with up-regulated TNFR1 expression was significantly higher (211.09 months, *p*-value=0.0366) than that of patients in the subgroup with normal expression (127.23 months), while the median survival of patients with down-regulated TNFR1 expression was 114.06 months (*p*-value=0.8783) (**Figure 5A/B**). On the contrary, patients with down-regulated expression of TNFR2 showed an inferior median overall survival 89.09 months when compared to the median survival of 127.23 months for patients with normal receptor expression (*p*-value=0.0376). The median survival of patients with upregulated TNFR2 expression was 244.91 months, but this was not statistical significant (*p*-value=0.146) when compared to the median survival of patients with normal receptor expression (**Figure 5C/D**).

**Figure 5.**
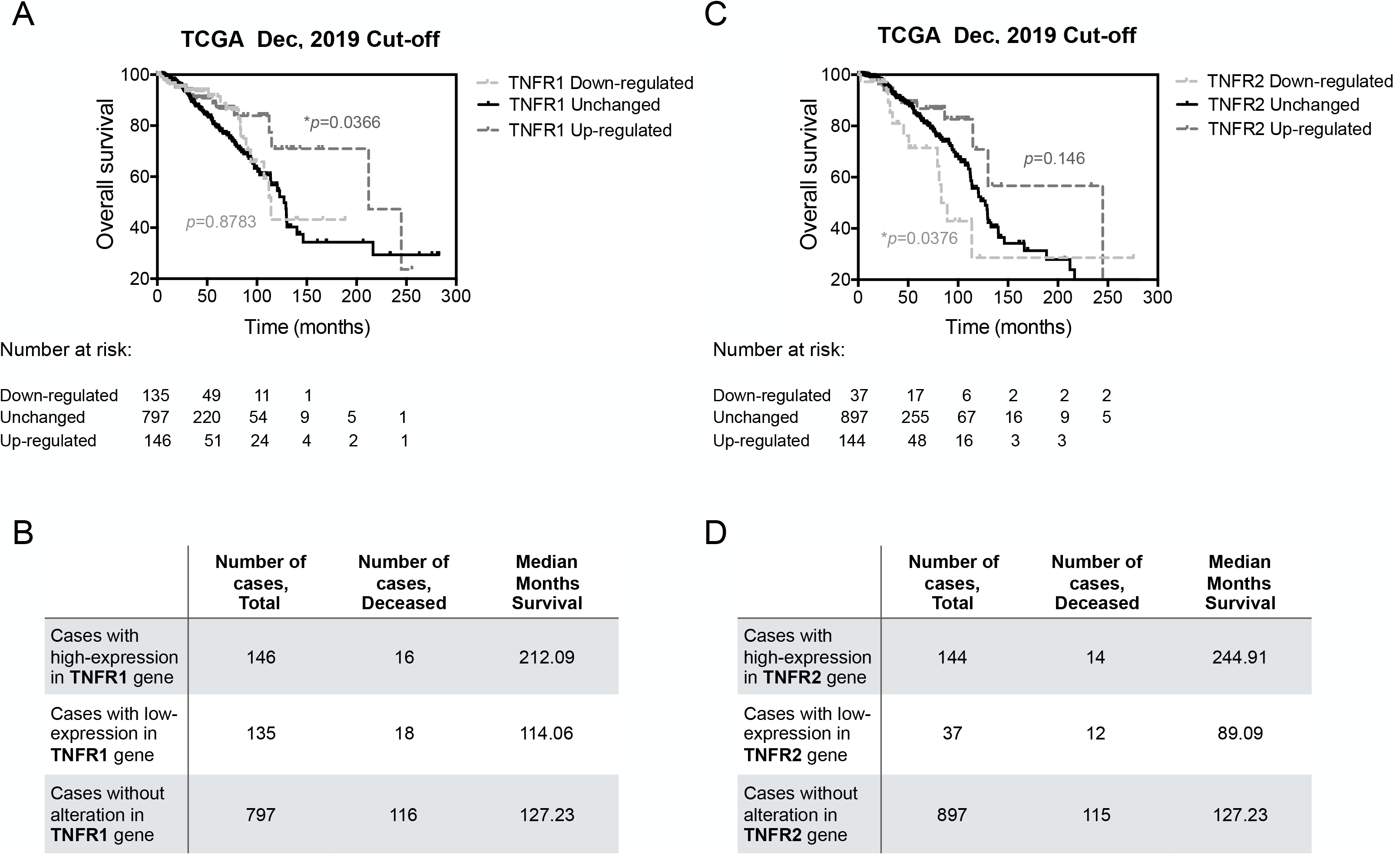
Overall survival evaluation of TNFR1 and TNFR2 signature in TCGA breast cancer patients. **(A)** The overall survival (OS) in breast cancer patients from TCGA stratified by TNFR1 mRNA expression. The light grey dotted line indicates patients with down-regulated expression of TNFR1, the dark grey dotted line represents the patients with up-regulated TNFR1 expression and the black line represents the unchanged population. OS in TNFR1 up-regulated subgroup was significantly better than that of the unchanged group (log rank test, *p*-value=0.0366). **(B)** The total case number, the deceased case number and the median survival of patients with down-regulated, unchanged or up-regulated TNFR1 expression**. (C)** OS in breast cancer patients from TCGA stratified by TNFR2 mRNA expression. The light grey dotted line indicates patients with down-regulated expression of TNFR2, the dark grey dotted line represents the patients with up-regulated TNFR2 expression and the black line represents the unchanged population. OS in TNFR2 down-regulated subgroup was significantly worse than that of the unchanged group (log rank test, *p*-value=0.0376). The number of patients at risk are listed below the Kaplan-Meier curves. **(D)** The total case number, the deceased case number and the median survival time of patients with down-regulated, unchanged, or up-regulated TNFR2 expression. *P* values were calculated using Log-rank (Mantel-Cox) test. * *p*-value<0.05.

## Discussion

Proinflammatory conditions are in large part driven by TNFα and have long been known to promote the development of malignancies including breast cancer (24, 25). In tumors, TNFα plays a critical role in proliferation, angiogenesis, invasion and metastasis via signaling through the TNFα/NF-κB axis (26). However, it has been unclear, which signaling events downstream of TNFα receptors affect tumorigenesis.

In this study we show that the homozygous knockout of either TNFR1 or TNFR2 alone increased the mammary epithelial stem cell numbers and led to accelerated prepubescent mammary gland development with hyperplastic ductal outgrowth and larger gland areas when compared to age-matched C57BL/6 animals. However, over the past 20 years of breeding p55 and p75 KO animals we have never observed an increase in spontaneous breast cancer in either of the TNFR knock-out strains, consistent with the resistance of their genetic C57BL/6 background strain against spontaneous breast cancer development (27). At 6-weeks of age, MMTV-Wnt1 x p75 KO animals had increased numbers of total, as well as M1 and M2 macrophages associated with mammary epithelial cells, thus implicating a role of the microenvironmental in the hyperplastic ductal outgrowth. Furthermore, in the litters of TNFR2 KO mice that were crossed with MMTV-Wnt1 transgenic animals that are prone to develop mammary tumors (18), the onset of breast cancer was significantly accelerated and the tumors showed a more aggressive phenotype.

Compared to less aggressive tumors developing in MMTV-Wnt1 animals, tumors in MMTV-Wnt1 x p75 KO animals showed elevated DNA-binding capacity of the NF-κB subunit p50, which indicated a preferential response of these tumors to TNF*α* and Interleukine-1 (IL-1) through NF-κB p50/p50 homodimers and the canonical NF-κB signaling pathway. In contrast, tumors in MMTV-Wnt1 x p55 KO animals showed elevated DNA-binding capacity of the NF-κB subunit p52, which signals through the non-canonical NF-κB pathway, does not respond to TNF*α*, and exclusively depends on IKK*α* but not IKKβ and IKKγ (28).

Consistent with an activation of the canonical NF-κB signaling pathway, tumor cell lines derived from MMTV-Wnt1 x p75 KO mammary tumors showed significantly upregulated levels of the pro-inflammatory cytokines IFN-γ, IL-1*α*, TNF*α*, IL-12 and the chemotaxis-related CXCL1 chemokine. Treatment of tumor cell lines derived from MMTV-Wnt1 and MMTV-Wnt1 x p55 KO breast tumors as well as the triple-negative human breast cancer line SUM159PT with exogenous TNF*α* led to a dose-dependent increase in mammosphere formation and also an increased plating efficacy in a colony-forming assays. On the contrary, tumor cell lines derived from MMTV-Wnt1 x p75 KO breast tumors did not respond to TNF*α*but showed higher baseline mammosphere formation comparable to that of MMTV-Wnt1 and MMTV-Wnt1 x p75 KO cell lines stimulated with high concentrations of TNF*α*, results consistent with an autocrine pro-inflammatory loop. It is noteworthy that when breast cancer-initiating cells (BCICs) were purged from SUM159PT cells as described previously (8), treatment with TNF*α* or radiation led to re-expression of the developmental transcription factors Oct4, Sox2 and Klf4 and that coincided with a phenotype conversion in some of the remaining non-BCICs into induced BCICs. The combined treatment with radiation and a high concentration of TNF*α*appeared to have an additive effect. These finding were in line with our previous report on radiation-induced phenotype conversion in breast cancer (7). Inflammatory breast cancer is a rare and aggressive form of the disease with worse survival than other types of breast cancer (29). Our observation of TNF*α*-induced phenotype conversion could be one aspect of the aggressive phenotype of inflammatory breast cancer.

Finally, we analyzed the overall survival of 1078 breast cancer patients in the TCGA Provisional dataset. Patients with upregulated TNFR1 expression showed significantly increased overall survival while patients with down-regulated TNFR2 expression had a significantly reduced overall survival, thus supporting the clinical significance of our experimental findings.

We are not the first to report an effect of TNF*α* on the mammary gland. TNF*α* has been shown to increase the growth of both normal and malignant mammary epithelial cells in experimental models (30, 31) and has been associated with tumorigenesis by affecting stem cell fate and inducing the transformation of normal stem cells (32, 33). TNF*α* exerts its activity by stimulation of its receptors TNFR1 and TNFR2. These two receptors trigger distinct and common signaling pathways that control cell apoptosis and survival (5). However, to our best knowledge the effects of imbalances in TNFR1 and TNFR2 signaling on mammary gland development and carcinogenesis have never been studied in literature. In our present study, we elucidated the role of TNFRs in mammary gland development and carcinogenesis using genetic mouse models. While expression of the MMTV-Wnt1 transgene in our model was restricted to the mammary epithelium, TNFRs were knocked out globally, suggesting that the effects observed were not only from changes in the mammary epithelium, the tumor cells derived from it or the surrounding microenvironment, but could have rather also accounted for systemic changes of immunity and inflammation. By testing the effects of TNF*α* on mammary epithelial cells and mammary tumor cells derived from our different mouse strains *in vitro* we were able to separate these effects, demonstrating a clear role for TNF*α* in the self-renewal of normal mammary epithelial stem cells as well as breast cancer-initiating cells.

Macrophages are important regulators of mammary gland developmental processes, especially during puberty stage when they are recruited to the neck region of the TEBs and guide the mammary gland ducts out-branching (34). Furthermore, macrophages are mediators of inflammation, play an indispensable role in modulating innate immune responses and interacting as tumor-associated macrophages (TAMs) with the surrounding microenvironment (35). However, the effects of both the resident macrophages and the TAMs on tumorigenesis or progression are still incompletely understood. In our present study, we did not perform longitudinal studies to address whether macrophages were responsible for increased ductal outgrowth and numbers of TEBs or if instead TNFR signaling imbalances in epithelial cells were the primary cause for this phenotype with macrophages secondarily associating with the ductal system. Future studies are warranted to address these outstanding questions.

We conclude that the roles of TNFR1 and TNFR2 in breast cancer development and progression are rather complex and go far beyond the pro-inflammatory properties of TNF*α*. It still needs to be determined if selective targeting of the receptors by e.g. specific TNFR2 agonists can modulate mammary carcinogenesis and disease progression.

## Acknowledgements

FP was supported by grants from the *National Cancer Institute* (R01CA161294, 5R01CA200234, P50CA211015) and a grant from the *National Institute of Allergy and Infectious Diseases* (AI067769).

